# Methylsulfonylmethane and sesame seed oil improve hyperlipidemia and polyunsaturated fatty acid metabolism in the *db/db* mouse

**DOI:** 10.1101/2020.12.30.424717

**Authors:** Cameron V. Fili, Ling Lin, Jonathan Chapman, David Hamilton, Charles R. Yates

## Abstract

**Background:** An estimated 463 million people worldwide suffer from diabetes, and that number is projected to rise significantly to 700 million by 2045. One of the hallmarks of type 2 diabetes and metabolic syndrome is alterations in the lipid profile and polyunsaturated fatty acid metabolism.

**Objective:** The objective of this study was to identify lipid alterations in leptin receptor deficient *db/db* mice, a type 2 diabetes model, to establish a baseline biological signature for the evaluation of natural products with purported lipid-altering activity.

**Methods:** Six-week old male *db/db* mice (n = 10/group) were randomized to the following groups: 1) diabetic control with no treatment, 2) methylsulfonylmethane (MSM) treatment (3.81 ± 0.33 g/kg), 3) sesame seed oil (SSO) treatment (23.54 ± 2.91 mg/kg), and 4) MSM and SSO combination treatment, receiving the same dosages of MSM and SSO as the single treatment groups. Eight-week old male C57BL6/J mice (n=10) were used as a non-diabetic control group.

**Results:** Serum triglycerides and total cholesterol were significantly increased in the *db/*db model compared to nondiabetic control, mimicking diabetes in people. High-density lipoprotein (HDL) was significantly increased in all *db/db* treatment groups, with the most significant effect in the MSM and SSO combination group, with a corresponding decrease in non-HDL cholesterol. Serum total polyunsaturated fatty acid (PUFA) levels were significantly reduced in diabetic mice compared with control mice. In contrast, treatment with SSO alone reversed this effect such that fed mice exhibited serum PUFA levels comparable to control mice.

**Conclusions:** Treatment of *db/db* mice with MSM and SSO improved commonly measured clinical parameters in serum lipid panels. The combination of MSM and SSO treatment’s effects on HDL and non-HDL cholesterol and fatty acid metabolism could lead to improved clinical outcomes in diabetics, such as reduced incidence of atherosclerosis and hepatic steatosis.

## Introduction

The International Diabetes Federation estimates there are at least 463 million people (1 in 11 adults aged 20-79 years) worldwide suffering from diabetes, and that number is projected to rise significantly to 700 million by 2045 [1]. One of the hallmarks of type 2 diabetes mellitus and related metabolic syndrome and obesity is dyslipidemia and impaired lipid metabolism [1, 2]. While most type 2 diabetic patients require pharmaceutical intervention such as insulin therapy and peroxisome proliferator-activated receptor (PPARγ) agonists, changes in the diet may also improve dyslipidemia and the diabetic condition [3]. An important contributor to the rise in the number of diabetic patients is the increasing rates of obesity due to the Modern Western Diet. In the past three decades, total fat and saturated fat intake as a percentage of total calories has continuously increased in Western diets, while the intake of omega-6 fatty acids increased and omega-3 fatty acids decreased. This shift has resulted in a large increase in the omega-6: omega-3 ratio from 1:1 during evolution to 20:1 today or even higher [4]. This high omega-6: omega-3 ratio is directly linked to chronic inflammatory diseases including obesity, metabolic syndrome, and type 2 diabetes mellitus [5].

Current anti-diabetic therapeutic strategies, e.g., insulin, metformin, and PPARγ agonists, are employed once an individual has already developed type 2 diabetes. A more recent approach to controlling the diabetes epidemic is to elucidate the mechanisms of pre-diabetes as a means to understand factors that contribute to conversion of pre-diabetes to diabetes. Pre-diabetes, or impaired glucose tolerance and insulin resistance prior to onset of type 2 diabetes, has been historically diagnosed based on glycemic criteria including hemoglobin A1c of 5.7-6.4%, fasting blood glucose of 100-125 mg/dL, and blood glucose of 140-199 mg/dL two hours post oral glucose tolerance test [1, 6]. Early detection of the pre-diabetic condition and insulin resistance (IR) can lead to intervention and prevention of the development of type 2 diabetes [1]. The recent development of tests such as the Quantose™ IR test, which measures biomarkers of insulin sensitivity other than blood glucose and A1c, can detect pre-diabetes even earlier in the disease process than these standard glycemic markers [7]. One biomarker included in the Quantose™ IR is linoleoylglycerophosphocholine (LGPC), also known as lysophosphatidylcholine-linoleic acid (LPC-LA, 18:2), which is a representative omega-6 polyunsaturated fatty acid (PUFA) that decreases in people with pre-diabetes [6]. Dietary supplements and foods that can improve this dyslipidemia found in the diabetic and pre-diabetic patient could lead to improved clinical outcomes; thus there is a growing need for research into lipid altering compounds.

The dietary ingredient sesamin, a phytonutrient of the lignans class found abundantly in sesame seed oil, is known to decrease formation of the pro-inflammatory omega-6 metabolite arachidonic acid in the liver by inhibiting the delta-5 desaturase activity. This effect results in enhanced liver formation of eicosapentaenoic acid (EPA) and docosahexaenoic acid (DHA) relative to arachidonic acid (AA) [8]. Sesamin’s impact on the PUFA balance is limited by the fact that the liver inefficiently delivers EPA and DHA to the systemic circulation and thus the tissues in the setting of diabetes. This occurs because a key contributor to PUFA release from the liver is the presence of methyl donors such as vitamin B12 and folic acid, which are significantly reduced in diabetes [9]. Methylsulfonylmethane, with the chemical formula (CH_3_)_2_SO_2_, also known as MSM, DMSO2, methyl sulfone, or dimethyl sulfone, is a naturally-derived sulfur compound found in many plants and food sources and a putative methyl donor [7, 8]. MSM provides remarkable defense against oxidative stress and inflammatory injuries [9, 10]. The objective of this research was to investigate whether supplementation with sesame seed oil (SSO), MSM, or the combination of SSO and MSM will improve the dyslipidemia seen in the diabetic condition, and thus reverse the pro-inflammatory environment found in an animal model of type 2 diabetes mellitus, the leptin receptor deficient *db/db* mouse.

## Materials and methods

### Animals and housing

Six-week old male BKS.Cg-*Dock7*^*m*^ +/+ *Lepr*^*db*^/J (hereafter referred to as *db/db*) mice were purchased from The Jackson Laboratory as a model for type 2 diabetes mellitus. This strain is leptin receptor deficient; therefore, leptin is unable to bind and produce satiety, resulting in development of obesity and type 2 diabetes mellitus. Eight-week old male C57BL6/J mice (n=10) were used as a non-diabetic control group. Nondiabetic mice (n=10) received no treatment, and diabetic mice were divided into four experimental groups (n=10 per group): 1) diabetic control with no treatment, 2) MSM treatment, 3) SSOtreatment, and 4) MSM and SSOcombination treatment. Upon arrival and prior to treatments, mice were placed in individually ventilated caging at a housing density of 5 mice per cage on corncob bedding (Shepherd Specialty Papers, Watertown, TN) with a 12:12 light: dark cycle. Chlorinated water was delivered in polysulfone bottles (Alternative Design, Siloam Springs, AR) ad libitum. Mice were fed Tekland 7912 rodent diet ad libitum (Envigo, Madison, WI). Animal work was performed in an Association for Assessment and Accreditation of Laboratory Animal Care International (AAALAC) accredited animal facility and approved by the University of Tennessee Health Science Center Institutional Animal Care and Use Committee (Protocol 17-095.0).

### Diabetes confirmation and treatments

One week after arrival, non-fasted blood glucose was measured to confirm hyperglycemia in the *db/db* mice. Blood glucose was measured by handheld glucometer (FreeStyle Lite) on a drop of blood obtained from the tail vein with the mouse in a plastic mouse restraining device. Mice with blood glucose measurements greater than 250 mg/dL were considered diabetic. Records were maintained of daily individual mouse weight, daily water and food intake per cage of five mice, and weekly individual mouse blood glucose levels. The criteria for early euthanasia was 20% weight loss from the start of experiment or hunched posture, however, none of the mice in this study reached these euthanasia criteria. After confirmation of diabetes, *db/db* mice then began treatment with either MSM alone, SSO alone, or a combination of MSM and SSO for four weeks prior to euthanasia via isoflurane overdose. MSM provided by Bergstrom Nutrition was dissolved in Millipore filtered water in a polysulfone bottle (Alternative Design, Siloam Springs, AR) at one percent v/v. Tekland 7912 feed was soaked in sesame seed oil (Sigma Aldrich) for five minutes, blotted dry with an absorbent paper towel, then stored in a sealed plastic container and fed ad libitum. Nondiabetic C57BL6/J mice and diabetic control *db/db* mice continued to receive unaltered 7912 diet and Millipore filtered water in polysulfone bottles for four weeks prior to euthanasia via isoflurane overdose.

All drug solutions were sterile filtered using a 0.22-micron filter prior to administration. Blood was collected via cardiocentesis under isoflurane anesthesia at euthanasia, and then centrifuged at 2500 rpm for fifteen minutes for serum separation. Serum was aliquoted for Rodent Lipid Profiles and fatty acid analysis via LC-MS/MS. Samples of 250 uL of serum per mouse were mailed on ice overnight to IDEXX BioAnalytics for Rodent Lipid Profile measurement, including cholesterol, high-density lipoprotein (HDL) cholesterol, and triglyceride levels. Non-HDL cholesterol was calculated by subtracting HDL cholesterol from total cholesterol. At euthanasia, tissues (eyes, brain, and liver) were harvested for fatty acid analysis using LC-MS/MS. Sections of liver were preserved in formalin, cryosectioned, and stained with Oil Red O for lipid content. Histology slides were read and scored numerically from 1 to 5 for degree of hepatocyte steatosis, or fatty change, by a blinded veterinary pathologist. The scores of 1 through 5 were none, mild, moderate, marked, and pathologic, respectively.

### LC-MS/MS parameters

LC-MS/MS analysis was performed following previous papers with minor modifications [11, 12]. The LC-MS/MS system comprised a Sciex (Framingham, MA) 5500 triple quadrupole mass spectrometer, equipped with a Turboionspray™ ionization interface in negative ion mode. The heated capillary temperature was 600°C, the sheath gas pressure was 50 psi, the auxiliary gas setting was 50 psi, and the heated vaporizer temperature was 350°C. The collision gas was argon at a pressure of 1.5 mTorr. The analytical data were processed using the software program Analyst (Version 1.3). Analyses were performed in the multiple reaction monitoring (MRM) mode, in negative ion mode. The collision energy (CE), collision cell exit potential (CXP), entrance potential (EP), and declustering potential (DP) were optimized for each compound to obtain optimum sensitivity.

A reverse-phase column (Analytical UPELCO C18; 2.1mm ID; 15mm length; 3 um particle size; 175A pore size) with a gradient elution of solvent A (5 mM ammonium formate in water, pH 4.0) and solvent B (5 mM ammonium formate in 95% (v/v) acetonitrile, pH 4.0) was used for chromatographic separation. Flowrate was at 0.5 ml/min. Solvent A was prepared by mixing 95 ml of Milli-Q (Millipore) water and 5 ml of 100 mM ammonium formate, then pH was adjusted to 4.0 with formic acid. Similarly, solvent B was prepared by mixing 95 ml acetonitrile and 5 ml of 100 mM ammonium formate, adjusting pH 4.0 with formic acid. The gradient started at 30% of mobile phase B, maintained 30% B for 1 minute and then followed by a linear gradient to 100% B from 1 to 8 minutes, and held at 100% B for 1 minute. Subsequently, the mobile phase was immediately returned to the initial condition 30% B and maintained for 1 minute until the end of the run. Column effluents were introduced into the mass spectrometer between 1 and 9 minutes after injections. The run time for a single sample was 11 minutes with most of the lipids eluting from 3 minutes to 5 minutes.

### Lipid extraction

Liver, brain, and eye tissues were weighed and placed into 1.5 ml siliconized sample tubes. 500 µl volumes of acidic methanol (pH 4.0) with 200 ng/ml deuterium-labeled arachidonic acid (AA-d_6_) were added for the internal standard. The obtained mixtures were homogenized for 1 minute by Bio-Gen PRO200 Homogenizer (with a 5 mm probe, 3,000 rpm, on ice). Mixtures were continuously homogenized for 10 minutes on ice in an ultrasonic bath. After an initial centrifugation at 1,000 g for 10 minutes at 4°C, the supernatants were further centrifuged at 13,500 g for 10 minutes at 4°C. The resulting supernatants were filtered with a 0.2 µm pore size and 4 mm inner diameter filters. Approximately 150 µl of filtrations were transferred to a 96-well plate for LC-MS/MS.

Serum was incubated at 37°C for 1 hour, allowed to stand at 4°C for 12 hours, and centrifuged at 1,500 g for 10 minutes at 25°C. The serum samples (30 µl each) were placed in 1.5 ml siliconized sample tubes, deproteinized by mixing with 150 µl acidic methanol (pH 4.0), and homogenized for 10 minutes on ice in an ultrasonic bath. After an initial centrifugation at 1,000 g for 10 minutes, the supernatants were further centrifuged at 13,500 g for 10 minutes at 4°C, as was done with the liver, brain, and eye samples. The resulting supernatants were filtered and 150 µl of filtrations were subjected to LC-MS/MS.

### Statistics

Mouse weights, blood glucose measurements, and Rodent Lipid Panels, were analyzed via two-way ANOVA with Tukey posthoc analysis for statistical significance. Results of the fatty acid analysis via LC-MS/MS were analyzed via Kruskal-Wallis H test for statistical significance. All data were analyzed in SPSS and considered significant if the p value was less than 0.05.

## Results

### Weight, water and food intake, and blood glucose

Mice gained weight appropriately according to their age throughout the study, and no mice reached early euthanasia criteria. There were no significant differences in diabetic mouse weights among the treatment groups at the end of treatment (Fig 1). Diabetic mice weighed significantly more than nondiabetic control mice (p<0.05), which was to be expected due to the *db/db* obese condition. The average daily intake of MSM in water was 4.49 ± 0.53 g/kg of body weight for the MSM treatment group and 3.81 ± 0.33 g/kg of body weight for the MSM + SSO treatment group. The average daily intake of the ingredient sesamin in sesame seed oil-soaked rodent feed was 23.54 ± 2.91 mg/kg of body weight for the SSO treatment group and 22.70 ± 2.57 mg/kg of body weight for the MSM + SSO treatment group. Blood glucose readings on day 0 and day 28 of treatment did not significantly differ within any treatment group, although the blood glucose of *db/db* mice on day 28 of treatment with MSM (326 ± 21 mg/dL) was mildly decreased compared with day 0 of treatment (375 ± 30 mg/dL).

**Fig 1.**
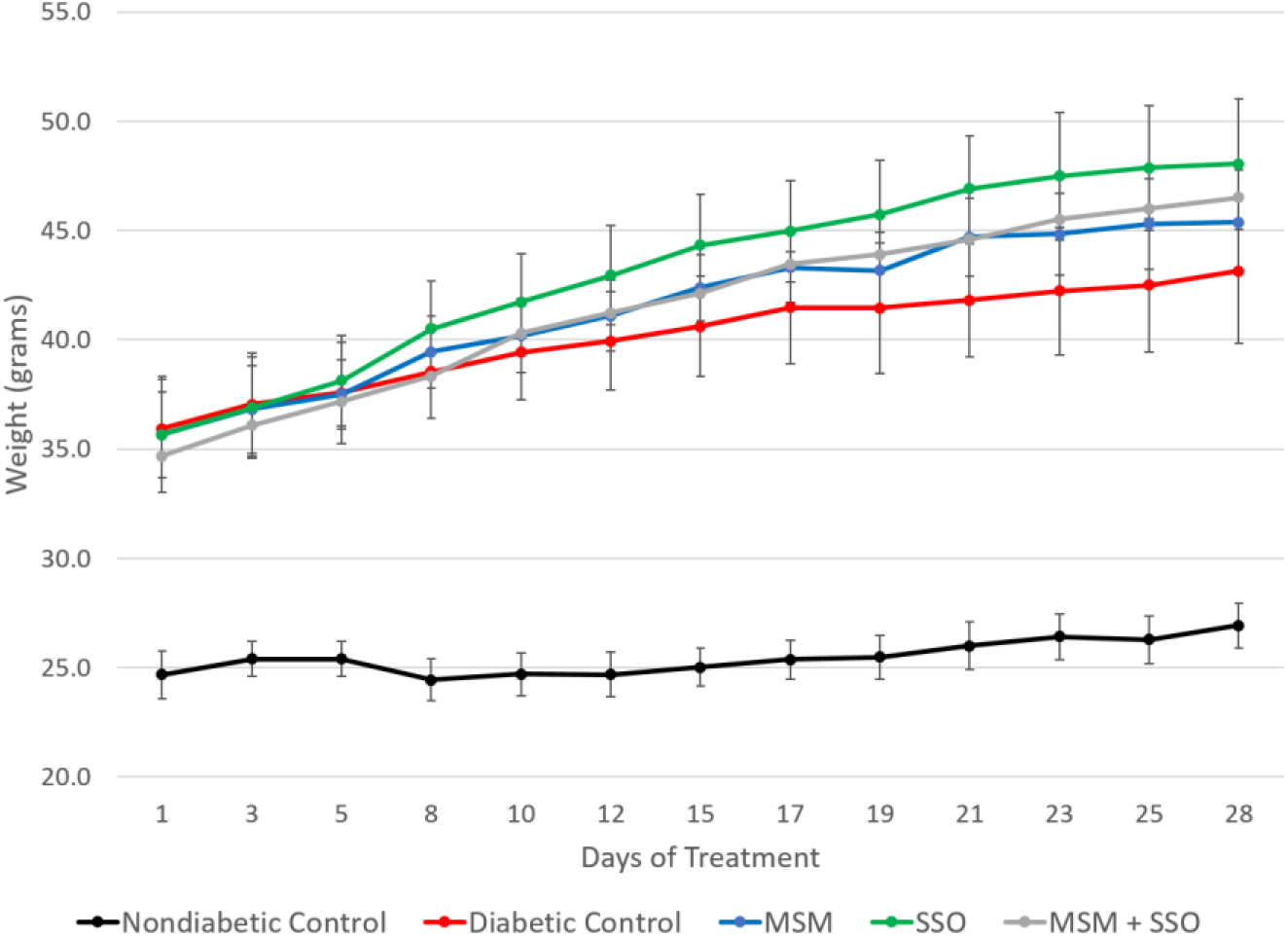
Type 2 Diabetic Mice Weights. Body weights of nondiabetic C57BL/6J mice and *db/db* mice over the 4-week treatment period.

### Rodent lipid panels

Results of the Rodent Lipid Panels received from IDEXX demonstrated significant differences between the nondiabetic control group and the diabetic groups. Serum cholesterol was significantly increased in the type 2 diabetic control and all type 2 diabetic treatment groups compared with the nondiabetic control group (Fig 2). Serum triglycerides were significantly increased in the diabetic control, SSO treated, and MSM plus SSO treated diabetic groups compared with the nondiabetic control group. In the MSM treated diabetic mice, serum triglycerides were not significantly different from the nondiabetic control group (Fig 3).

**Fig 2.**
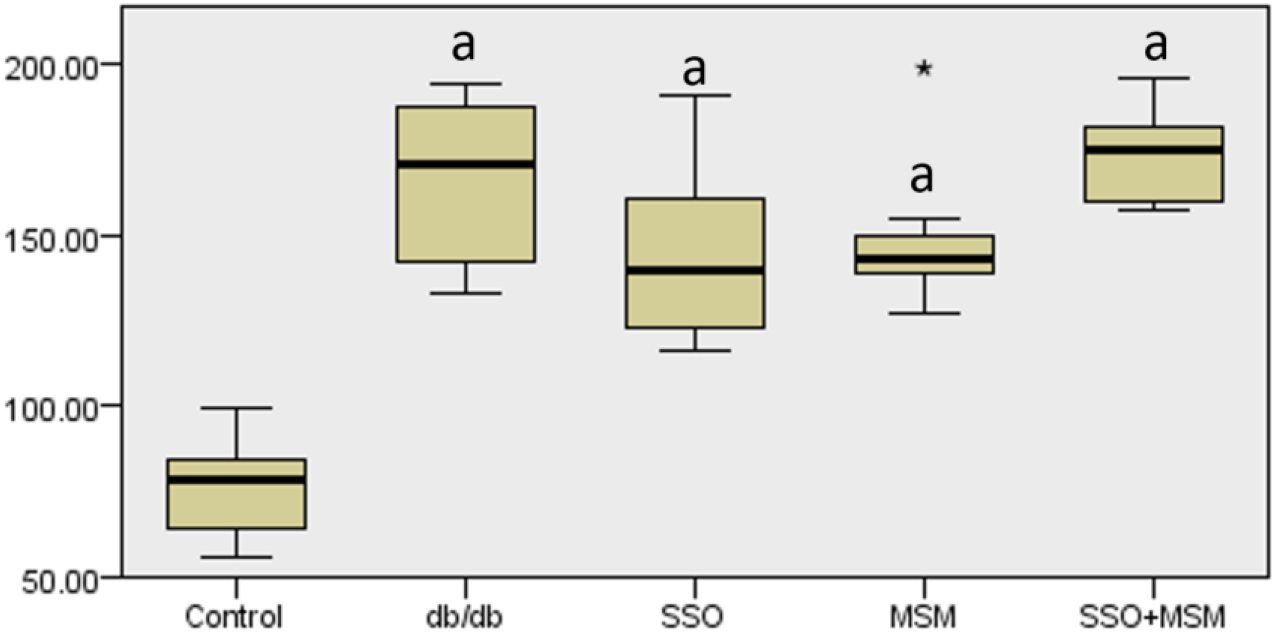
Total Cholesterol. Total cholesterol levels of C57BL/6J mice (non-diabetic control) and *db/db* mice after the 4-week treatment period, where a denotes p<0.05 compared with nondiabetic control and * denotes one outlier measurement.

**Fig 3.**
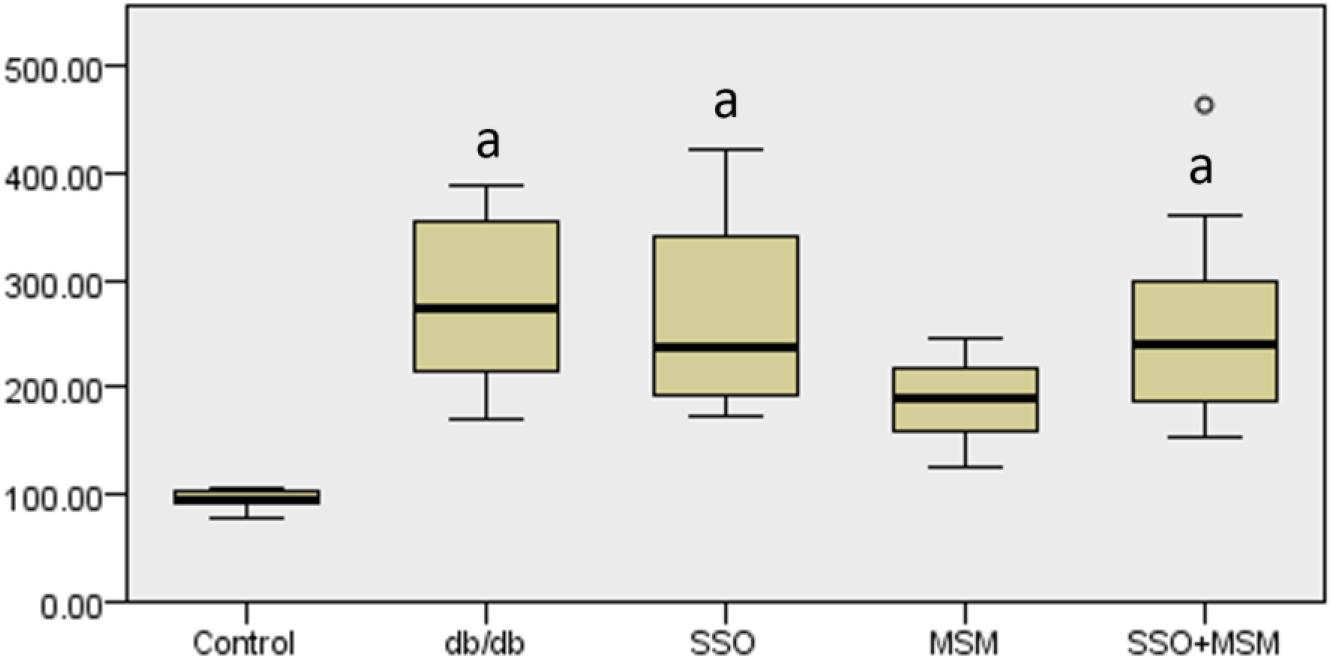
Triglycerides. Triglyceride levels of C57BL/6J mice (non-diabetic control) and *db/db* mice after the 4-week treatment period, where a denotes p<0.05 compared with nondiabetic control and ° denotes one outlier measurement.

Serum HDL was significantly increased in all diabetic groups compared with the nondiabetic control group. Diabetic mice treated with MSM, SSO, or a combination of the two had significantly increased HDL levels compared with diabetic control mice. Additionally, the HDL levels of diabetic mice receiving treatment with MSM/SSO combination were significantly increased compared with treatment with either MSM alone or SSO alone (Fig 4). Serum non-HDL cholesterol levels were significantly increased in all diabetic groups compared with the nondiabetic control. There was a decreasing trend in non-HDL cholesterol levels in all diabetic mice receiving treatment with either MSM, SSO, or combination of the two, although the differences were not statistically significant (Fig 5).

**Fig 4.**
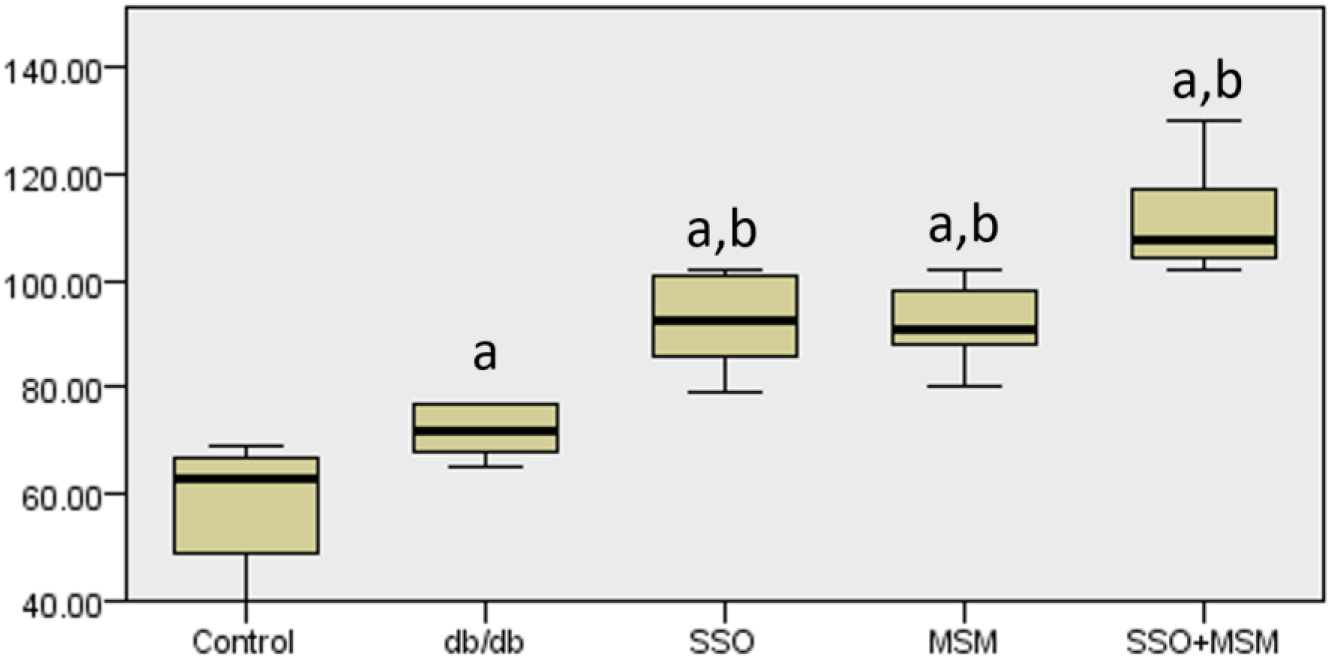
HDL Cholesterol. HDL cholesterol levels of C57BL/6J mice (non-diabetic control) and *db/db* mice after 4-week treatment period, where a denotes p<0.05 compared with nondiabetic control and b denotes p<0.05 compared with diabetic control.

**Fig 5.**
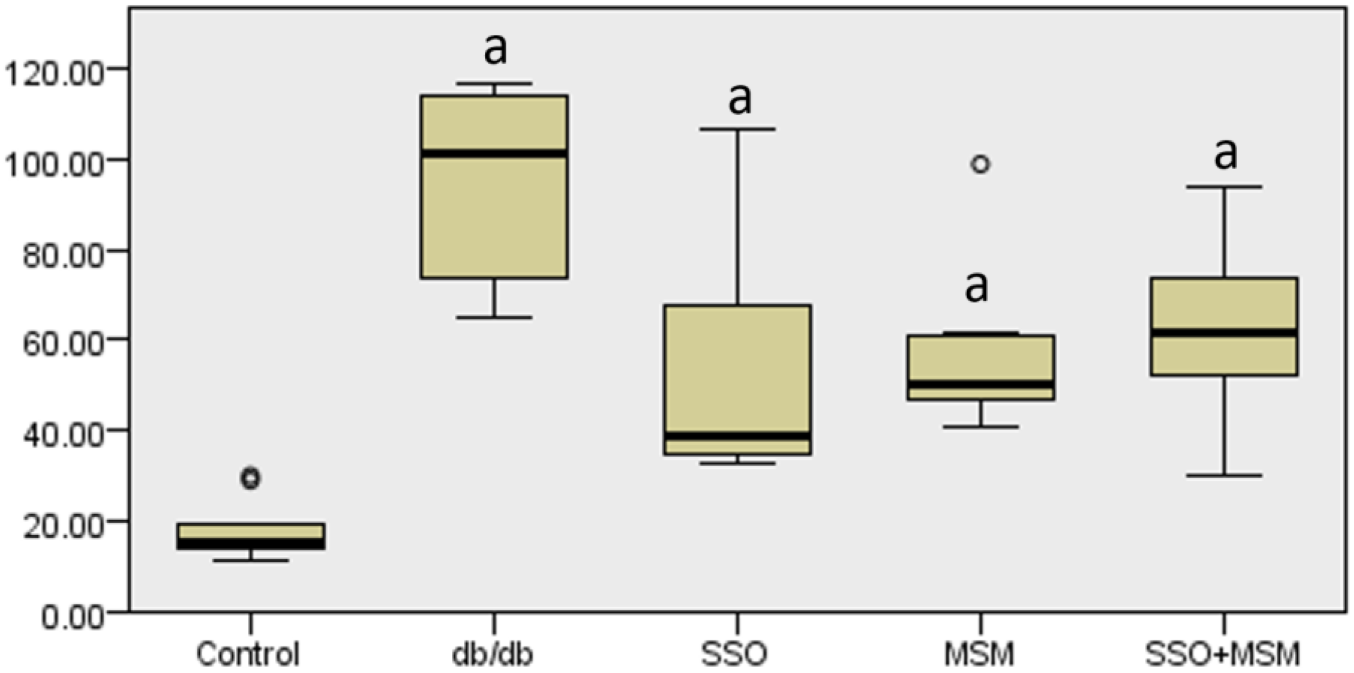
Non-HDL Cholesterol. Non-HDL cholesterol levels of C57BL/6J mice (non-diabetic control) and *db/db* mice after 4-week treatment period, where a denotes p<0.05 compared with nondiabetic control and ° denotes outlier measurements.

### Liver histology

In the diabetic mice, liver histology revealed variable microvesicular steatosis with intrahepatic variability and increased red-staining lipid found in the centrilobular regions around the central veins of the liver. There was intragroup variability and no significant differences among treatment groups (Fig 6A-D). All liver histology slides were scored either a 1 for appearing normal or a 2 for mild hepatic steatosis, but there was no trend among the groups, and therefore, no statistics were performed on the liver histology scores.

**Fig 6.**
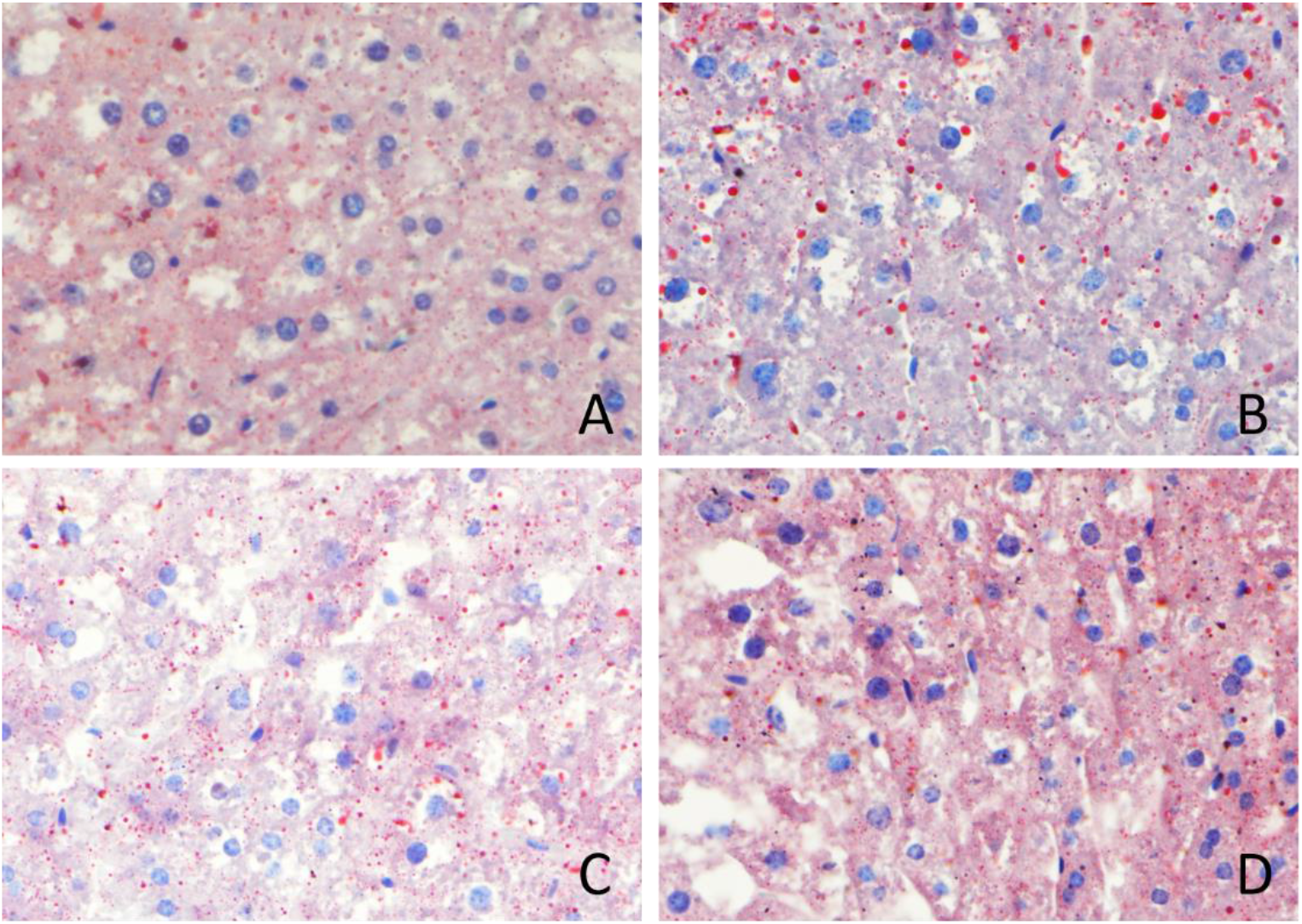
Liver Histology. Representative liver histology stained with Oil Red O from diabetic control group (A), diabetic mice treated with MSM (B), diabetic mice treated with SSO (C), and diabetic mice treated with both MSM and SSO (D).

### Free Fatty Acids

#### Serum

In the serum, linoleic acid (LA) levels were significantly increased in *db/db* mice treated with SSO or MSM compared with nondiabetic and diabetic control mice (Table 1). Dihomo-gamma-linolenic acid (DGLA) and arachidonic acid (AA) levels were significantly decreased in diabetic control mice compared with nondiabetic control mice, and this effect was reversed in mice treated with either SSO or MSM alone, where these levels were significantly increased compared with *db/db* control mice and not significantly different from nondiabetic control mice. In diabetic control and diabetic mice treated with MSM/SSO combination therapy, alpha linolenic acid (ALA) levels were significantly increased compared with nondiabetic controls.

**Table 1.** Serum Free Fatty Acids.

| Free fatty acids | Non-diabetic Control<br>(Mean ± SD) | Diabetic Control<br>(Mean ± SD) | SSO Treatment<br>(Mean ± SD) | MSM Treatment<br>(Mean ± SD) | MSM + SSO Treatment<br>(Mean ± SD) |
| --- | --- | --- | --- | --- | --- |
| AA | 102 ± 8.37 | 52 ± 17<sup>a</sup> | 117 ± 7.46<sup>b</sup> | 113 ± 14.6<sup>b</sup> | 79 ± 19 |
| DGLA | 1.1 ± 0.3 | 0.13 ± 0.06<sup>a</sup> | 1.5 ± 0.1<sup>b</sup> | 1.7 ± 0.5<sup>b</sup> | 0.17 ± 0.06<sup>a</sup> |
| LA | 4.2 ± 0.6 | 3.8 ± 1.1 | 6.3 ± 1.0<sup>ab</sup> | 5.9 ± 1.2<sup>ab</sup> | 5.4 ± 1.1 |
| DHA | 157 ± 12.7 | 100 ± 29.7<sup>a</sup> | 148 ± 13.2<sup>b</sup> | 156 ± 18.4<sup>b</sup> | 107 ± 23.4<sup>a</sup> |
| EPA | 30 ± 7.5 | 34.4 ± 14.0 | 41 ± 7.2 | 97 ± 12.8<sup>ab</sup> | 17 ± 7.8 |
| ALA | 0.6 ± 0.2 | 1.7 ± 0.5<sup>a</sup> | 0.89 ± 0.21<sup>b</sup> | 1.0 ± 0.3 | 1.8 ± 0.4<sup>a</sup> |
| NPD1 | 0.03 ± 0.01 | 0.04 ± 0.01 | 0.05 ± 0.02 | 0.03 ± 0.02 | 0.04 ± 0.01 |
| PGE2 | 0.10 ± 0.01 | 0.16 ± 0.07<sup>a</sup> | 0.13 ± 0.01<sup>a</sup> | 0.12 ± 0.03 | 0.12 ± 0.03 |
| total omega-3 | 187 ± 19.6 | 136 ± 42.7 | 189 ± 18.1 | 254 ± 30.5<sup>b</sup> | 126 ± 30.7<sup>a</sup> |
| total omega-6 | 107 ± 8.56 | 56 ± 18<sup>a</sup> | 15 ± 7.94<sup>b</sup> | 122 ± 15.9<sup>b</sup> | 84 ± 19 |
| Total PUFAs | 295 ± 25.3 | 193 ± 60.0 | 314 ± 25.2<sup>a</sup> | 376 ± 45.1<sup>a</sup> | 210 ± 48 |
| omega-6/omega-3 | 0.6 ± 0.1 | 0.41 ± 0.05<sup>a</sup> | 0.66 ± 0.04<sup>b</sup> | 0.48 ± 0.03 | 0.7 ± 0.1<sup>b</sup> |
| AA/DHA | 0.7 ± 0.1 | 0.52 ± 0.06 | 0.8 ± 0.1<sup>ab</sup> | 0.73 ± 0.04<sup>b</sup> | 0.7 ± 0.1<sup>b</sup> |
| AA/EPA | 3.7 ± 1.1 | 1.6 ± 0.3<sup>a</sup> | 2.9 ± 0.4<sup>a</sup> | 1.2 ± 0.1<sup>a</sup> | 5.2 ± 1.5<sup>b</sup> |

Serum free fatty acid levels of C57BL/6J mice (non-diabetic control) and *db/db* mice after 4-week treatment period, measured as a ratio with the internal standard, AA-d<sub>6</sub>.
<sup>a</sup>p<0.05 compared with nondiabetic control
<sup>b</sup>p<0.05 compared with diabetic control
<sup>ab</sup>p<0.05 compared with nondiabetic control and with diabetic control

In diabetic mice treated with SSO alone, ALA levels were returned to levels that were not significantly different from nondiabetic control and were significantly decreased compared with diabetic control mice. EPA levels were significantly increased in diabetic mice treated with MSM alone compared with both nondiabetic control and diabetic control mice. DHA levels were significantly decreased in diabetic control mice and diabetic mice treated with MSM/SSO combination therapy compared with nondiabetic control, but this effect was reversed in diabetic mice treated with SSO alone or MSM alone, whose DHA levels were significantly increased compared with diabetic control mice and not significantly different from nondiabetic control mice. Levels of neuroprotectin D1 (NPD1), a mediator derived from DHA that protects eye and brain from oxidative stress, was not significantly different among groups. Prostaglandin E2 (PGE2), a proinflammatory mediator derived from AA, was significantly increased in diabetic control mice and mice treated with SSO compared with nondiabetic control mice, but PGE2 was not significantly different in MSM treated or MSM/SSO treated mice compared with nondiabetic control mice.

#### Liver

The livers of diabetic mice treated with SSO alone and MSM/SSO combination therapy had significantly increased levels of LA compared with nondiabetic controls (Table 2). ALA levels were significantly increased in diabetic control mice and diabetic mice treated with either SSO alone or MSM alone compared with nondiabetic control mice. This effect was reversed in diabetic mice treated with MSM and SSO combined, and the ALA levels were not significantly different from nondiabetic control mice. EPA levels were significantly increased in diabetic control mice and diabetic mice treated with MSM alone compared with nondiabetic control mice but were significantly decreased in those treated with SSO alone compared with diabetic control. Mice receiving MSM/SSO combination therapy had levels of EPA comparable to those seen in nondiabetic control mice, and the two groups were not significantly different. There were no significant differences regarding AA,DHA, NPD1, or PGE2 levels among groups.

**Table 2.**
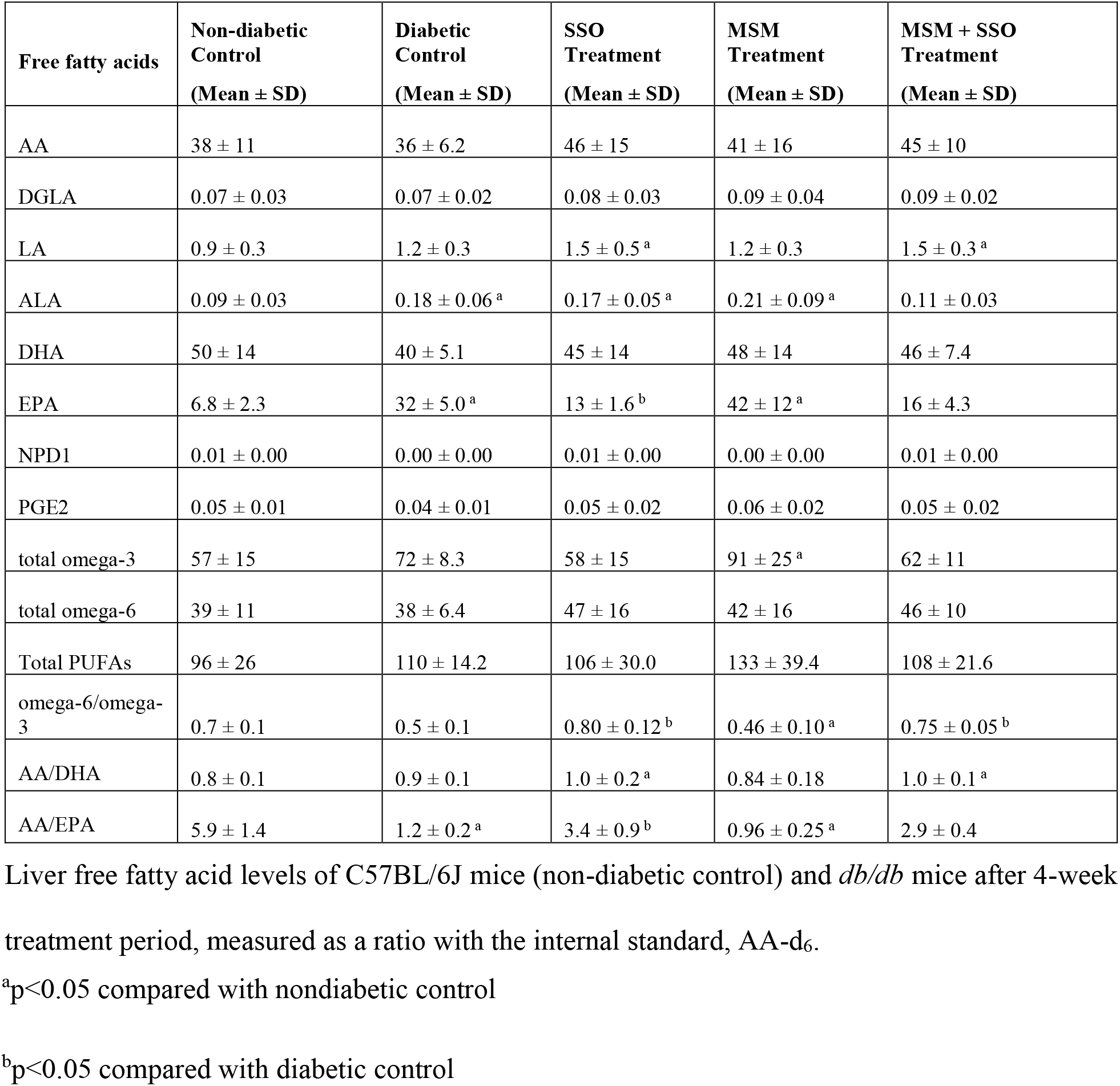
Liver Free Fatty Acids.

| Free fatty acids | Non-diabetic Control<br>(Mean ± SD) | Diabetic Control<br>(Mean ± SD) | SSO Treatment<br>(Mean ± SD) | MSM Treatment<br>(Mean ± SD) | MSM + SSO Treatment<br>(Mean ± SD) |
| --- | --- | --- | --- | --- | --- |
| AA | 38 ± 11 | 36 ± 6.2 | 46 ± 15 | 41 ± 16 | 45 ± 10 |
| DGLA | 0.07 ± 0.03 | 0.07 ± 0.02 | 0.08 ± 0.03 | 0.09 ± 0.04 | 0.09 ± 0.02 |
| LA | 0.9 ± 0.3 | 1.2 ± 0.3 | 1.5 ± 0.5<sup>a</sup> | 1.2 ± 0.3 | 1.5 ± 0.3<sup>a</sup> |
| ALA | 0.09 ± 0.03 | 0.18 ± 0.06<sup>a</sup> | 0.17 ± 0.05<sup>a</sup> | 0.21 ± 0.09<sup>a</sup> | 0.11 ± 0.03 |
| DHA | 50 ± 14 | 40 ± 5.1 | 45 ± 14 | 48 ± 14 | 46 ± 7.4 |
| EPA | 6.8 ± 2.3 | 32 ± 5.0<sup>a</sup> | 13 ± 1.6<sup>b</sup> | 42 ± 12<sup>a</sup> | 16 ± 4.3 |
| NPD1 | 0.01 ± 0.00 | 0.00 ± 0.00 | 0.01 ± 0.00 | 0.00 ± 0.00 | 0.01 ± 0.00 |
| PGE2 | 0.05 ± 0.01 | 0.04 ± 0.01 | 0.05 ± 0.02 | 0.06 ± 0.02 | 0.05 ± 0.02 |
| total omega-3 | 57 ± 15 | 72 ± 8.3 | 58 ± 15 | 91 ± 25<sup>a</sup> | 62 ± 11 |
| total omega-6 | 39 ± 11 | 38 ± 6.4 | 47 ± 16 | 42 ± 16 | 46 ± 10 |
| Total PUFAs | 96 ± 26 | 110 ± 14.2 | 106 ± 30.0 | 133 ± 39.4 | 108 ± 21.6 |
| omega-6/omega-3 | 0.7 ± 0.1 | 0.5 ± 0.1 | 0.80 ± 0.12<sup>b</sup> | 0.46 ± 0.10<sup>a</sup> | 0.75 ± 0.05<sup>b</sup> |
| AA/DHA | 0.8 ± 0.1 | 0.9 ± 0.1 | 1.0 ± 0.2<sup>a</sup> | 0.84 ± 0.18 | 1.0 ± 0.1<sup>a</sup> |
| AA/EPA | 5.9 ± 1.4 | 1.2 ± 0.2<sup>a</sup> | 3.4 ± 0.9<sup>b</sup> | 0.96 ± 0.25<sup>a</sup> | 2.9 ± 0.4 |

Liver free fatty acid levels of C57BL/6J mice (non-diabetic control) and *db/db* mice after 4-week treatment period, measured as a ratio with the internal standard, AA-d<sub>6</sub>.
<sup>a</sup>p<0.05 compared with nondiabetic control
<sup>b</sup>p<0.05 compared with diabetic control

#### Brain

In the brain, LA levels were significantly decreased in diabetic mice treated with MSM alone compared with diabetic control mice (Table 3). Diabetic control mice and diabetic mice treated with MSM/SSO combination therapy had significantly increased DGLA levels compared with nondiabetic controls. This effect was reversed in diabetic mice treated with MSM alone, whose DGLA levels were significantly decreased compared with diabetic control mice and were not significantly different from nondiabetic control mice. AA levels in diabetic mice treated with MSM alone were significantly increased compared with both nondiabetic and diabetic control mice. Diabetic control mice and diabetic mice treated with MSM alone had significantly increased levels of EPA. This effect was reversed in diabetic mice treated with SSO or MSM/SSO combination therapy, with EPA levels that were not significantly different from nondiabetic control mice and were significantly decreased compared with diabetic control mice. DHA levels were significantly increased in diabetic mice treated with MSM alone compared with nondiabetic control mice. There was no significant difference in ALA, NPD1, or PGE2 levels among the groups.

**Table 3.** Brain Free Fatty Acids.

| Free fatty acids | Non-diabetic Control<br>(Mean ± SD) | Diabetic Control<br>(Mean ± SD) | SSO Treatment<br>(Mean ± SD) | MSM Treatment<br>(Mean ± SD) | MSM + SSO Treatment<br>(Mean ± SD) |
| --- | --- | --- | --- | --- | --- |
| AA | 1,025 ± 195.9 | 1,163 ± 314.1 | 1,254 ± 451.9 | 1,503 ± 205.1 <sup>ab</sup> | 1,358 ± 323.9 |
| LA | 9.7 ± 2.2 | 12 ± 2.0 | 14 ± 3.8 | 8.7 ± 1.8 <sup>b</sup> | 13 ± 2.5 |
| DGLA | 7.2 ± 2.9 | 15 ± 4.0 <sup>a</sup> | 11 ± 3.1 | 10 ± 1.4 <sup>b</sup> | 12 ± 2.4 <sup>a</sup> |
| DHA | 1,039 ± 223.5 | 1,181 ± 354.5 | 1,150 ± 340.7 | 1,490 ± 226.1 <sup>a</sup> | 1,365 ± 312.9 |
| EPA | 98 ± 21 | 150 ± 29.3 <sup>a</sup> | 82 ± 18 <sup>b</sup> | 199 ± 33.6 <sup>a</sup> | 105 ± 28.6 <sup>b</sup> |
| ALA | 1.0 ± 0.8 | 1.5 ± 0.5 | 1.2 ± 0.4 | 1.1 ± 0.4 <sup>b</sup> | 1.3 ± 0.5 |
| NPD1 | 0.5 ± 0.8 | 0.4 ± 0.3 | 0.23 ± 0.15 | 0.2 ± 0.1 | 0.3 ± 0.2 |
| PGE2 | 1.7 ± 0.7 | 1.9 ± 0.3 | 1.7 ± 0.3 | 2.0 ± 0.3 | 2.1 ± 0.4 |
| total omega-3 | 1,138 ± 222.4 | 1,333 ± 369.7 | 1,233 ± 355.4 | 1,690 ± 243.3 <sup>a</sup> | 1,471 ± 330.9 |
| total omega-6 | 1,042 ± 197.0 | 1,189 ± 312.1 | 1,278 ± 452.8 | 1,522 ± 204.5 <sup>a</sup> | 1,382 ± 324.0 |
| total PUFAs | 2,180 ± 402.2 | 2,521 ± 675.7 | 2,511 ± 775.1 | 3,212 ± 446.2 <sup>a</sup> | 2,853 ± 645.7 |
| omega-6/omega-3 | 0.9 ± 0.1 | 0.90 ± 0.07 | 1.0 ± 0.2 <sup>b</sup> | 0.90 ± 0.02 | 0.9 ± 0.1 |
| AA/DHA | 1.0 ± 0.1 | 1.00 ± 0.09 | 1.1 ± 0.2 | 1.01 ± 0.03 | 1.0 ± 0.1 |
| AA/EPA | 11 ± 4.1 | 7.8 ± 2.0 | 15 ± 4.0 <sup>b</sup> | 7.7 ± 1.0 | 14 ± 3.8 <sup>b</sup> |

Brain free fatty acid levels of C57BL/6J mice (non-diabetic control) and *db/db* mice after 4-week treatment period, measured as a ratio with the internal standard, AA-d<sub>6</sub>.
<sup>a</sup>p<0.05 compared with nondiabetic control
<sup>b</sup>p<0.05 compared with diabetic control
<sup>ab</sup>p<0.05 compared with nondiabetic control and with diabetic control

#### Eye

In the eye, the levels of NPD1, LA, and ALA were significantly decreased in all diabetic mice regardless of treatment compared with nondiabetic control mice (Table 4). DGLA and AA were significantly increased in diabetic mice receiving MSM, SSO, or MSM/SSO combination therapy compared with nondiabetic control mice. Diabetic mice receiving SSO or MSM/SSO combination treatment had significantly decreased EPA levels compared with diabetic control mice and were not significantly different from nondiabetic control mice. DHA was significantly increased in diabetic mice treated with MSM alone or in combination with SSO compared with nondiabetic control mice. There were no significant differences in PGE2 levels among the groups.

**Table 4.**
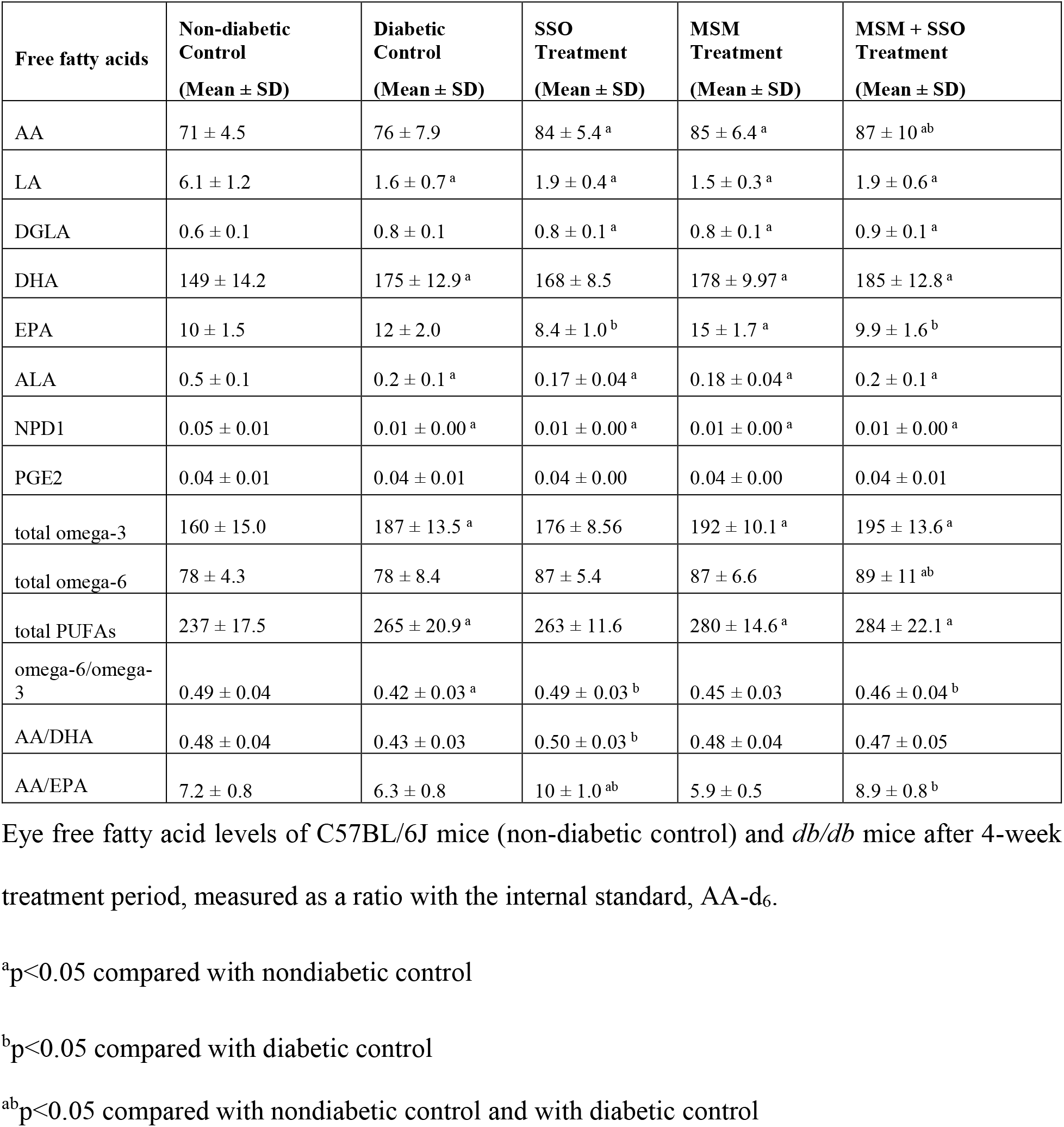
Eye Free Fatty Acids.

## Discussion

Dyslipidemia and impaired lysophospholipid metabolism are hallmarks of type 2 diabetes mellitus and related metabolic syndrome and obesity [1]. Diet and exercise, along with administration of cholesterol-lowering pharmaceuticals such as statins, are recognized as standard interventions to control hyperlipidemia. In recent years, however, there has been increasing interest and compelling evidence to support the adjuvant use of natural products to help control hyperlipidemia [13]. Our goal was to identify lipid alterations in *db/db* mice, a murine model of type 2 diabetes, for the purpose of establishing a baseline biological signature for the subsequent evaluation of natural products with purported lipid-altering activity. In this article, we utilized mass spectrometry-based lipidomics to investigate diabetes-associated changes in fatty acid metabolism and the extent to which the dietary ingredients sesame seed oil and/or methylsulfonylmethane impacted biomarkers of metabolic syndrome.

Sesamin, the primary constituent lignan found in SSO, exerts potent lipid-lowering effects by targeting key enzymes fatty acid synthesis (e.g., delta-5 desaturase), fatty acid oxidation (e.g., PPARα), and cholesterol metabolism (e.g., 3-hydroxy-3-methyl-glutaryl-CoA reductase) [14]. For example, low-density lipoprotein (LDL) receptor deficient mice fed a diet of SSO were found to have reduced plasma total cholesterol, TG, very low-density lipoprotein cholesterol (VLDL), and LDL cholesterol, while high-density lipoprotein (HDL) was significantly increased compared with atherosclerotic diet-fed animals [15]. Similarly, we found that diabetic mice fed SSO alone showed decreased non-HDL cholesterol (LDL + VLDL) accompanied by an increase in HDL (Fig 4 and 5), an effect that has been well recognized in human patients. Sesame seed oil supplementation has been shown to increase HDL cholesterol with a corresponding decrease in total cholesterol, triglycerides, and LDL cholesterol in hypercholesterolemic patients [16] and in rheumatoid arthritis patients, as well as reduce body weight, body mass index, and systolic blood pressure [17]. Similarly, Sankar et al. found that supplementation with 35 grams of sesame seed oil daily for 60 days significantly reduces triglycerides, total cholesterol, and LDL cholesterol while significantly increasing HDL cholesterol in type 2 diabetics [18] and hypertensive patients [19]. In postmenopausal women supplemented with 50 grams of sesame powder daily for five weeks, the same decrease in total cholesterol and LDL:HDL cholesterol was seen, with a corresponding increase in antioxidant status [20]. Our results affirm these findings in the type 2 diabetic *db/db* mouse.

Regarding fatty acid metabolism (Fig 7), serum total polyunsaturated fatty acid (PUFA) levels were significantly reduced in diabetic mice compared to control mice (Table 1). This finding is consistent with literature data demonstrating that in diabetics there is a reduction in the activity of delta-5 desaturase, the rate-limiting step in PUFA metabolism [21]. In contrast, feeding of only SSO reversed this effect such that fed mice exhibited serum PUFA levels comparable to control mice. SSO contains high amounts of omega-6 PUFAs such as linoleic acid [22], which partially explains the effect of SSO on omega-6 PUFA levels. However, since SSO is not an appreciable source of omega-3 PUFAs, dietary PUFAs in SSO cannot entirely account for the increase in total PUFAs, *viz*., omega-6 plus omega-3 PUFAs. A clue to an alternative mechanism to explain SSO’s effect on total PUFA levels may be found in its effect on the delta-5 desaturase index (AA:DGLA), which represents a measure of PUFA metabolism where a higher ratio reflects greater delta-5 desaturase activity. The delta-5 desaturase activity in diabetic mice (∼500) was higher than either control (∼100) or SSO only fed mice (∼100). Paradoxically, it appears that total serum PUFAs are reduced in diabetic mice compared to SSO-only fed mice despite delta-5 desaturase being five times greater in diabetic mice. Regardless of the mechanism, reduced PUFA availability can significantly impact lysophospholipid metabolism.

**Fig 7.**
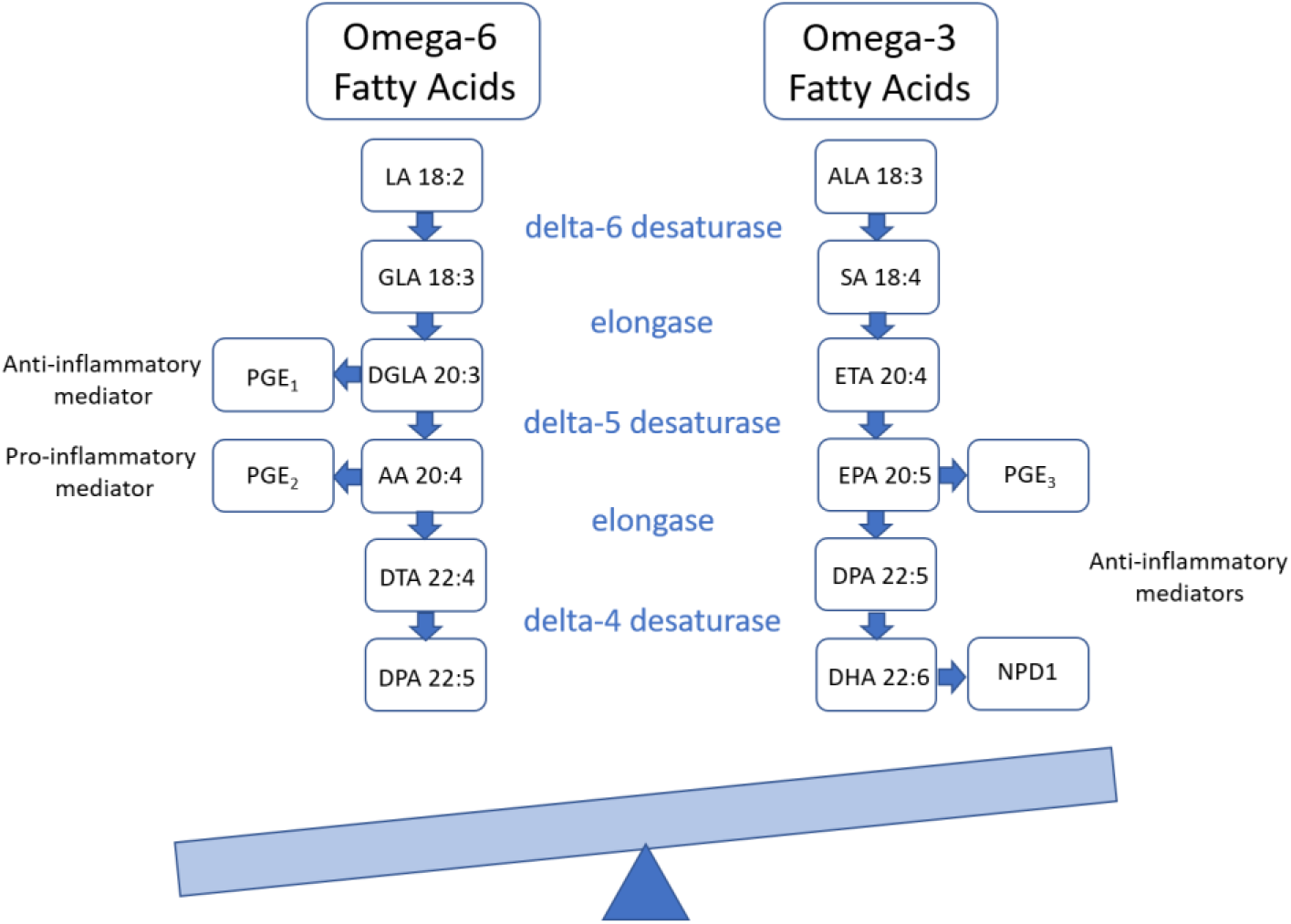
Fatty Acid Metabolism. This schematic demonstrates the metabolism of omega-6 and omega-3 fatty acids into pro-inflammatory and anti-inflammatory mediators, a balance which is disrupted in diabetes and metabolic syndrome.

Oral administration of methylsulfonylmethane to *db/db* mice results in a dose-dependent decrease in liver cholesterol, triglycerides, lipogenesis-dependent gene expression (e.g., fatty acid synthase), and fatty oxidation[8]. Since MSM contains two methyl groups, the authors speculated that MSM may serve as a methyl donor to ameliorate hepatic steatosis. Building on this hypothesis, we sought to demonstrate that MSM, acting as a methyl donor, could mitigate diabetes-related hyperlipidemia and aberrant PUFA metabolism. Similar to SSO, administration of MSM alone led to decreased non-HDL cholesterol (LDL + VLDL) and increased HDL (Fig 4 and 5).

The human genes for delta-5 (*FADS1*) and delta-6 (*FADS2*) desaturase enzymes are situated head-to-head (5’-5’) in a cluster configuration on chromosome 11 (11q12.2) [23]. This arrangement affords a common promoter for *FADS1* and *FADS2* that contains a potential “enhancer signature” comprising methylation loci located within a 5’ regulatory region [23]. Methylation probe studies demonstrated an inverse relationship between methylation proportion and both delta-5 (AA/DGLA) and delta-6 (GLA/LA) desaturase activity [23]. These data could potentially explain the association between methyl donor deficiency and reduced delta-5 desaturase activity in diabetics [21]. As mentioned above, our data demonstrate that delta-5 desaturase activity is greatly enhanced in *db/db* mice compared to control or MSM alone treated mice (Table 1). These data are consistent with the notion that reduced methyl donor availability in diabetic mice is associated with increased delta-5 desaturase activity and that administration of the putative methyl donor MSM significantly decreases delta-5 desaturase activity. Future studies examining the serum and liver methyl donor status (e.g., choline, betaine, etc.) in *db/db* mice are needed to test this hypothesis.

In conclusion, treatment of *db/db* mice with MSM and SSO improved commonly measured clinical parameters in serum lipid panels. Serum triglycerides and total cholesterol were significantly increased in the *db/db* model compared to nondiabetic control, mimicking the diabetic condition in people. Most notably, HDL-cholesterol was significantly increased in all *db/db* treatment groups, with the most significant treatment effect in the MSM + SSO combination group. Because the HDL-cholesterol level is increasing with treatment, the ratio of LDL:HDL is in turn decreasing. This change in the LDL:HDL ratio due to combination MSM and SSO treatment could lead to improved clinical outcomes in people such as a reduced incidence of atherosclerosis.

## Acknowledgements

The authors would like to acknowledge the Laboratory Animal Care Unit’s husbandry and cagewash technicians at the University of Tennessee Health Science Center for providing excellent daily care for their diabetic mice.

## Notes

### Competing Interest Statement

The authors have declared no competing interest.

### Summary of Updates

Methods, Results, and Discussion have been revised along with Tables and Figures

